# Computational and experimental evidence to the permeability of withanolides across the normal cell membrane

**DOI:** 10.1101/802645

**Authors:** R. Wadhwa, N. S. Yadav, S. P Katiyar, T. Yaguchi, C. Lee, H. Ahn, C-O. Yun, S. C Kaul, D. Sundar

**Affiliations:** AIST, Tokyo; Department of Biochemical Engineering & Biotechnology, DAILAB, Indian Institute of Technology (IIT) Delhi, Hauz Khas, New Delhi-110016; DAILAB, DBT-AIST International Center for Translational and Environmental Research (DAICENTER), National Institute of Advanced Industrial Science & Technology (AIST), Tsukuba - 305 8565, Japan

**Keywords:** Permeability, Withaferin-A, Withanone, POPC membrane, human normal and cancer cells

## Abstract

Poor bioavailability due to the inability to cross the cell membrane is one of the major reasons for the failure of a drug in the clinical trials. We have used molecular dynamics simulations to predict the membrane permeability of natural drugs - withanolides (withaferin-A and withanone) that have similar structures but remarkably differ in their cytotoxicity. We found that withaferin-A, but not withanone, could proficiently transverse through the model membrane. The free energy profiles obtained were in accordance with the physico-chemical properties of the investigated drug molecules. It was observed that the polar head group of the bilayer exhibits high resistance for the passage of withanone as compared to withaferin-A, while the interior of the membrane behaves similarly for both withanolides. The solvation analysis revealed that the high solvation of terminal O5 oxygen of withaferin-A was the major driving force. The impact of the favorable interaction of terminal oxygen (O5) of withaferin-A with the phosphate of the membrane led to its smooth passage across the bilayer. The computational predictions were validated by raising and recruiting unique antibodies that react to withaferin-A and withanone. Further, the time-lapsed analyses of control and treated human normal and cancer cells, demonstrated proficient permeation of withaferin-A, but not withanone, through normal cells. These data strongly validated our computational method for predicting permeability and hence bioavailability of candidate compounds in the drug development process.

**Statement of significance:** What determines the bioavailability of a drug? Does the ability to cross cell membrane determine this? A combined simulation/experimental study of the permeability of two natural drugs - withanolides (Wi-A and Wi-N) across the cell membrane was conducted. In the computational portion of the study, steered MD simulations were performed to investigate the propensity of the two molecules to permeate across the cell. It is found that Wi-A proceeds relatively simply across the cell compared to Wi-N. This trend was found to be consistent with experiment. This work is an important step towards understanding the molecular basis of permeability of natural drug molecules.

## Introduction

Drug design and development is a multidimensional and extensive regimen. Many potent drugs fail in later stages of trials leading to huge loss of time and resources. Inappropriate pharmacokinetics of molecules is one of the major causes of such failure. These pharmacokinetic properties generally include low bioavailability, short elimination half-life and variable behavior due to genetic or environmental factors. Especially for orally administered drugs, bioavailability is very critical for proper absorption that largely depends on their solubility and membrane permeability.

In eukaryotic systems, both active and passive transport modes are available for the transport of a molecule through a lipid membrane (1). Active transport requires ATP to transport the molecule across a membrane, while passive transport involves diffusion of molecule across the membrane without any external assistance or energy input. However, the rate of passive diffusion across a bilayer is proportional to the partition coefficient between membrane and external medium, diffusion coefficient of the compound through the membrane, and concentration gradient of compounds across the bilayer. Further, the important physico-chemical properties responsible for the process of membrane binding and diffusion are lipophilicity, molecular weight and polarity.

Studies on characterization and kinetics of drug permeability have shown that majority of molecules get absorbed in the body via passive diffusion for which a drug must penetrate the apical membrane. Thus, passive transport across the apical cell membrane represents an essential step of drug design, bioavailability and efficacy. Membrane penetration potential analysis of candidate drugs becomes important for elimination of molecules that may fail later due to poor bioavailability or chose the ones with specificity for only the diseased cells.

Withanolides (C_28_-steroidal lactones), a class of secondary metabolites from Solanaceae plant family are long known to possess anti-cancer (2–7), anti-stress, anti-neurodegenerative (8), and anti-microbial (9) activities. Withaferin-A (Wi-A) extracted from Ashwagandha (*Withania somnifera*) has been studied the most amongst all withanolides that share the same core structure (10). Interestingly, despite having quite similar structures, Wi-A and Withanone (Wi-N) differ remarkably in their activities. Wi-A is a C5, C6 epoxy compound carrying hydroxyl groups on C4 and C27, while Wi-N is a C6, C7 epoxy compound having hydroxyl groups on C5 and C17 (Fig. 1). Wi-N, although less potent as compared to Wi-A in cytotoxic assays for cancer cells, has been earlier shown to be safe for normal cells (11–14). Various factors such as molecular size, structural conformations, type and position of the functional groups affect the interactions of withanolides with their biological targets. Permeability across cell membrane can also be anticipated to be the reason behind this differential activity.

**Fig. 1:**
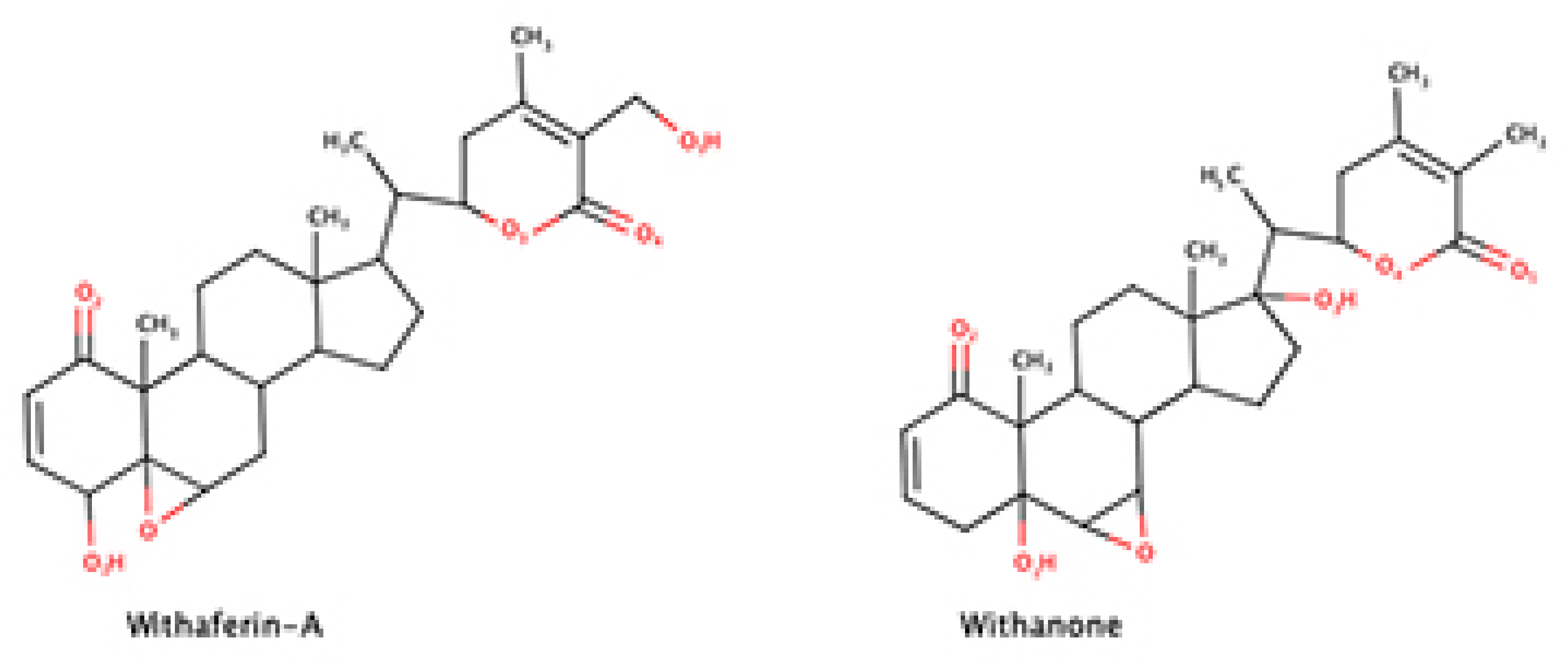
The 2D pictorial representation of the anti-cancer drug candidates.

Though experimental (15–17) and theoretical studies (18–20) have been carried out in the past for investigating drug permeability, Molecular Dynamics Simulations (MDS) is now widely used for the estimation of partition of a drug in a membrane (21, 22). Since *in vivo* studies of intestinal membrane permeability are expensive and may prove potentially harmful for the volunteers, the *in vitro* models such as partitioning in isotropic system (23, 24), transport across artificial membranes (25) and transport across cultured epithelial cell monolayer (26, 27) have been the valuable tools. However, these models do not take into account the molecular properties of membrane that play an important role in governing the permeability of a solute. Computational methods, on the other hand, hold remarkable potential to investigate the structure-permeability relationships. Recently, full atom computational models of membranes such as POPC (1-palmitoyl-2-oleoyl-sn-glycero-3-phosphocholine), POPS (1-palmitoyl, 2-oleoyl-sn-glycero-3-phosphoserine) *etc*. have been used to study the interactions of small molecules with the membrane and their permeability characteristics (28, 29). These membrane models exhibit the structural and dynamic features of membrane bilayer, which have been investigated and verified by MDS.

Cholesterol is one of the major components of eukaryotic cell membrane, and is also reported to have a large effect on the membrane permeability. Generally solute permeability across the high cholesterol content is low (30–32). Eriksson and Eriksson by measuring the potential of mean force (PMF) barriers of photodynamic drugs, hypericin and its derivatives have also shown that, cholesterol significantly influences the permeation of these molecules (33). They showed that the calculated rate of permeation is lower at high cholesterol content due to increase in PMF barrier. Nonetheless, in simulations, mixed membrane model allowing the mixing of cholesterol molecules with POPC membrane has been reported to mimic the real cell membrane system in a much better way (34). Such membrane models have been used successfully to predict the permeability of drugs such as ibuprofen, cimitedine, six-β-blocker, aliphatic amine, and carboxylic acid drugs using the MDS (35, 36).

The partition of a drug could be either predicted by long nano-second classical MDS, or by the Potential of Mean Force (PMF) methods (37, 38). Using MDS on various fluorescent azaaromatic probes on water-membrane, it has been earlier shown that the probe localization is determined by the electrostatic dipole-dipole and van der Waals interactions (39). Further, it was also shown that 2,6 Bis (1H-Benzimidazole-2-yl) pyridine (BBP) preferred to locate around 15-16Å from the DPPC (1,2dipalmityol-sn-glycero-3-phosphatidycholine) membrane center (37). A MDS study of 17 amino acids in lipid bilayer showed that the partitioning of charged and polar side chain amino acids were accompanied by water defects (40). Nonetheless, the solute size, polarizability and free energy of transfer from water to membrane also affected the transfer of solute from water into membrane.

In this study, we have elucidated the mechanism of permeation of two natural drug molecules (Wi-A and Wi-N) through POPC bilayer using computer simulations, which were further validated by recruiting unique withanolide-recognizing antibodies in cell-based assays. In-spite of having very little difference in their structures, Wi-A molecules successfully crossed the bilayer with low free energy associated with it, while the passage of Wi-N was associated with high free energy barrier. We present here a system that would be useful for predicting permeation and bioavailability of drug compounds in drug development process.

## Methods

### Lipid Membrane Simulations

Three MD simulations, (1) POPC membrane (2) POPC membrane having Withaferin-A (3) POPC membrane with Withanone, were carried out. To include the complexity of a real cell membrane and be fully miscible at the simulated conditions, an equilibrated POPC membrane from a 2µs long equilibration simulation was used (41). The bilayer membrane model consisted of 70 POPC and 35 randomly cholesterol molecules per layer or leaflet. All simulations were run using TIP3P water models (42) and amber Lipid14 force field parameter (43). It has been earlier shown that Lipid14 is a good choice for molecule permeability calculations (44), all the structures were converted to the Lipid14 naming convention by using charmmlipid2amber.py script, available with Amber Tools v18. The prepared models were then completely solvated, by 7499 TIP3P water molecules. To attain the charge neutrality, 19 Na^+^ and 19 Cl^−^ ions were added making the salt concentration of ∼0.15 M in each simulation box.

### Equilibration and Production procedure

The solvated membrane in the simulation box was then equilibrated using a slightly modified version of the multi-step protocol (45). Briefly the procedure consisted of altering cycles of steepest descent (1000 steps) and conjugate gradient energy minimization (1000 steps) followed by position restrained MD with strong harmonic restraints on all non-hydrogen atoms of the bilayer membrane and solvent. In successive steps the positional restraints were gradually relaxed. The detailed description of the minimization and equilibration protocol can be found in supplementary material. All the energy minimization and MD simulations were run using the SANDER modules of the AMBER16 program suite (46). Production simulations were started from the velocities and coordinates obtained from the last equilibration step. The production simulations for all the systems were run for 600 ns in NPT ensemble using GPU accelerated version of the PMEMD program of AMBER simulation suite (47, 48). The temperature and pressure of the systems were kept restrained to 310 K above the crystalline fluid/liquid phase transition temperature and 1 atm respectively using the Berendsen’s coupling scheme (*τ*_p_ = *τ*_t_ = 1 ps) (49). All bonds involving hydrogen atoms were constrained using the SHAKE algorithm. Short range electrostatics were calculated up to 10 Å and the Particle Mesh Ewald (50) algorithm was used to calculate the long-range electrostatics. An integration time step of 2 fs was used and structures were saved at every picosecond resulting in around 6,00,000 structures.

#### Potential of Mean Force

The potential of mean force (PMF) free energy profiles for the partitioning of drug compounds were calculated using umbrella-sampling simulations.

#### Umbrella Sampling

The reaction coordinates was defined in the Z direction between the phosphorous atom of lipid membrane and the atoms of drug molecules. For both permeants, 30 windows (upper leaflet) spaced 1 Å apart were used with a biasing harmonic constraint of 1.5 kcal/mol/Å^2^ was used. Each window was run for 30 ns, totaling ∼1 *μs* of sampling per permeant. Further, to calculate the PMF, the biased distributions were reweighted using Weighted Histogram Model (WHAM) (51, 52).

#### Diffusion and Resistance Calculations

Theoretically, the permeability coefficient of a molecule can be computed from an atomistic simulation-based PMF approach, via the inhomogeneous solubility diffusion permeability model (53). Here, the permeation is divided into a three-step process – (a) involving the partitioning of a permeant from the aqueous phase on the side of leaflet, (b) diffusion across the bilayer and (c) partitioning from the other side of bilayer into aqueous phase.

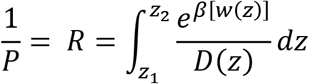

where, *R*(*z*) is the resistivity of every “slice” of the membrane at position “z”. *w*(*z*) is the PMF, *D*(*z*) is the local position specific diffusion coefficient, which can be calculated by the Hummer’s positional autocorrelation extension of Wolf-Roux estimator (54). A detailed discussion on the computation of local diffusion coefficient *D*(*z*) has been reviewed elsewhere (55–57).

#### Area per lipid (APL)

The area of a side of the system at any particular state was calculated by measuring the distance between farthest water molecules. The distance between the two farthest water molecules in x-plane gave the length of *a,* and the distance between the two farthest water molecules in y-plane gave the length of *b*. The total area was calculated as *a* * *b*. The total number of lipid molecules (70 molecules) in one leaflet was constant. Therefore, dividing the area by the total number of POPC molecules gave the area per lipid (APL) for that side as given below:

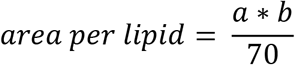

#### Order Parameter

The ordering of lipid acyl chains was determined by the calculation of order parameter S_CD_. This quantity can be directly compared to experimental S_CD_ values obtained by ^2^H NMR or ^1^H-^13^C NMR. Since S_CD_ is also a measure of relative orientation of the C-D bonds with respect to the bilayer normal, it can be calculated according to the following equation:

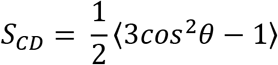

Where *θ* is the angle between the bilayer normal and vector joining C-D (actually C-H in the current simulation), and < > represents an ensemble average.

#### Generation of anti-Withaferin A and anti-Withanone antibodies

Wi-A and Wi-N were extracted from dried Ashwagandha leaves as described in a previous study (12). Mixture of Wi-A/Wi-N (1:1) along with Freund’s adjuvant was used as antigen to immunize the mice for monoclonal antibody generation. The resulting clones (∼30) were screened with affinity ELISA and one clone L7C3-6 (WiNA Ab) was isolated that reacted to Wi-A and Wi-N in fixed cells. The clone was established as hybridoma. Affinity purified antibody was used for this study.

#### Cell culture and treatments

Human normal (TIG-3, MRC5 and WI38) and cancer (breast carcinoma-MCF-7, melanoma-G361 and osteosarcoma-U2OS) were obtained from Japanese Collection of Research Bioresources (JCRB, Japan). The cells were authenticated by the source. Cells were frozen in −80°C and LN_2_ in multiple vials and were cultured in Dulbecco’s modified Eagle’s medium (DMEM; Gibco BRL, Grand Island, NY, USA) and treated either with Wi-A or Wi-N at about 60% confluency. Internalization of Wi-A and Wi-N was detected by immunostaining with the anti-WiNA Ab raised in our laboratory. The treated cells were also immuno-stained with a variety of other antibodies that included anti-γH2AX (Millipore #07-627), anti-ATR1 (Abcam, #ab4471), anti-CHK1 Cell Signaling #2345S), anti-p53 (Santa Cruz. #sc-126) and anti-CARF (58).

## Results and Discussion

### Membrane Architecture

In order to validate out simulation protocol, few parameters like area per lipid (APL), electron density and Lipid order were calculated in control as well as in presence of Wi-A and Wi-N molecules, which were compared to the experimentally observed values.

#### Area Per Lipid (APL)

The compactness of a biomembrane is measured by area per lipid. Addition of a permeant alter the density of a membrane. For the liquid-crystalline phase of POPC membrane, a number of APL values have been reported, depending upon the temperature: 68 Å^2^ at 303K (59), 62 Å^2^ at 323K, 66 Å^2^ at 310K (60), 63 Å^2^ at 310K (61), 62 Å^2^ at 310K (62). In our MD simulations, the APL for the control POPC membrane was ∼60.22 ± 0.06 Å (Fig. 2A), which increased slightly to 60.27 ± 0.055 Å^2^ and 60.28 ± 0.06 Å^2^ in the presence of Wi-A and Wi-N respectively (Fig. 2B and C). The little increment in APL in the presence of drug candidates indicates a small deformation in membrane architecture. Further the discrepancies, between the calculated and the experimental values of APL could be attributed to the presence of cholesterol molecules in the biomembrane. Using ^13^C-NMR spectroscopy on POPC membrane, it has been earlier reported that addition of 50% cholesterol caused ordering of the POPC membranes, by losing the entropy (63).

**Fig. 2:**
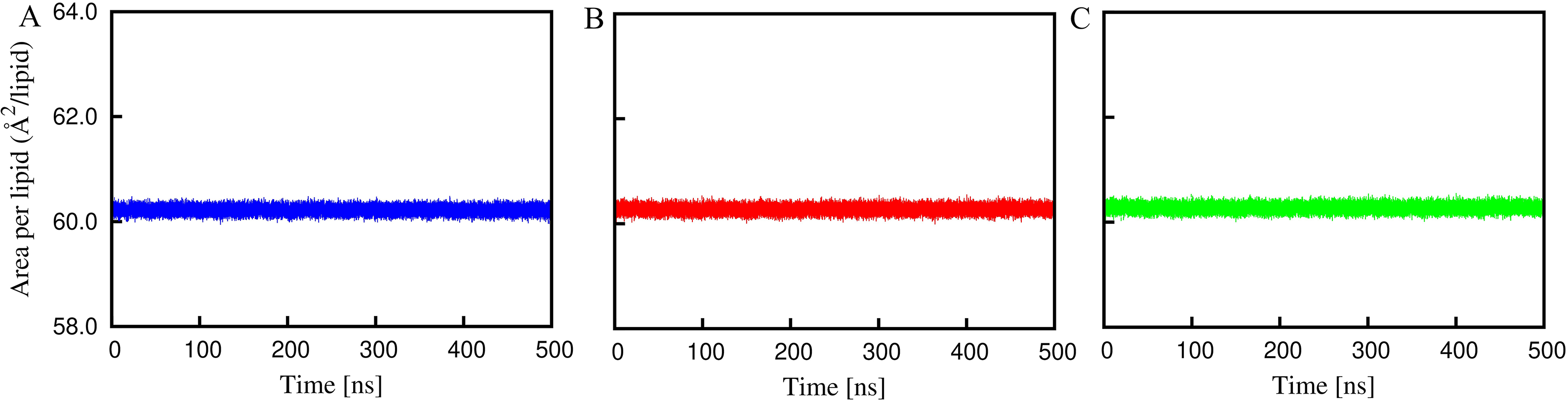
Lipid area variation of POPC membrane (A) in presence of Withaferin-A (B) and Withanone (C).

#### Electron Density Profile

Generally, thickness of the bilayer is determined by the electron density profile measured by the X-ray scattering of liquid-crystalline membranes (59, 61, 62). The electron density profiles (EDP) were calculated by assuming an electron charge equal to the atomic number minus the atomic partial charge, located at the center of each atom. The density profile for POPC bilayer in control and as well as in the presence of permeants are shown in Fig. 3. The electron density profile has been further decomposed into contributions from other groups such as - water, choline (CHOL), phosphate (PO_4_), glycerol, carbonyl (COO), methylene (CH_2_), unsaturated CH=CH and terminal methyl’s (CH_3_). Three distinct domains are evident from the profile shown in Fig. 3. The flat region between ∼20 Å to 30 Å represents the aqueous phase, unaltered from the bulk water, corresponds to the interface region, containing lipid head-groups and water. The drop in the density represents the inner part of the bilayer, where the alkyl chains of the phospholipid reside. The plan of the lipid density corresponds to the middle of the bilayer, where the ends of the tails of both lipid layers meet.

**Fig. 3:**
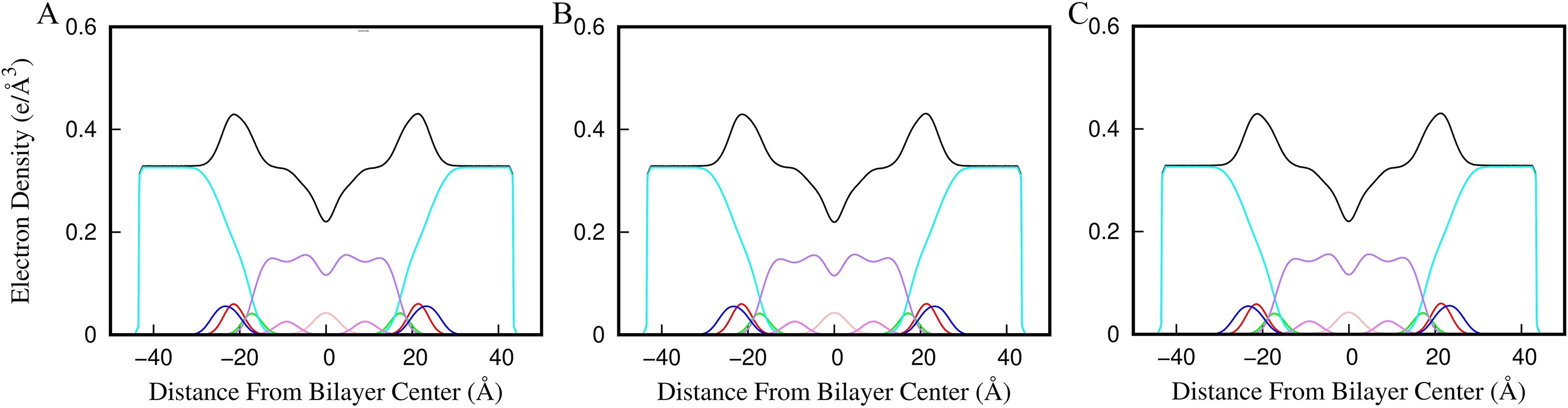
Electron density of control POPC (A) and POPC membrane in presence of Withaferin-A (B) and Withanone (C) molecule. The profiles of different membrane components are plotted where the black color corresponds to the total electron density variation. Water density is shown in cyan. The lipid atoms are in different colors – phosphate (red), carbonyl (green), choline (dark blue), double bond (magenta), methyl (pink) and methane (light blue).

In each case, the electron density profiles are symmetrical, almost overlapping, with water penetrating up to the carbonyl groups, leaving the terminal methyl groups dehydrated in agreement to the experimental results. The simulated electron density profile indicates that the parameters chosen for the MD simulations reproduces the membrane geometry at a good level.

#### Order Parameter (S_CD_)

The order parameters of POPC membrane acyl chain in presence of Wi-A and Wi-N are shown in Fig. 4. The calculated order values follow the similar trend as obtained by the previous simulation and experimental values (64, 65). The carbon-2 atoms of the sn-1 and sn-2 chains displays different order parameters owing to different alignment of the acyl chains in the region. Experimentally, it has been observed that the S_CD_ of C-D bonds near the head group in the sn-1 chains are greater than the sn-2 chains (66). Similar behavior was observed for the calculated lipid systems. The unsaturated chain of POPC shows a drop at the carbon 9 and 10 due to double bond. Further, the high S_CD_ value of sn-1 chain indicates a high order of acyl chain, as compared to oleyl chain of POPC. The plot reveals that for the sn-1 chain (i) a maximum S_CD_ of 0.2 near the aqueous phase, (ii) the S_CD_ is slightly high for the first five segments as a plateau and (iii) from the sixth carbon, the value gradually decreases up to ∼0.07. In case of sn-2 oleyl chain, (i) a maximum S_CD_ of ∼0.19 near the aqueous phase, (ii) a distinguishing drop at C2^’^ to 0.15 and (iii) a characteristic minima ∼0.02 at carbon 9 and 10, owing to the double bond of the oleyl chain.

**Fig. 4:**
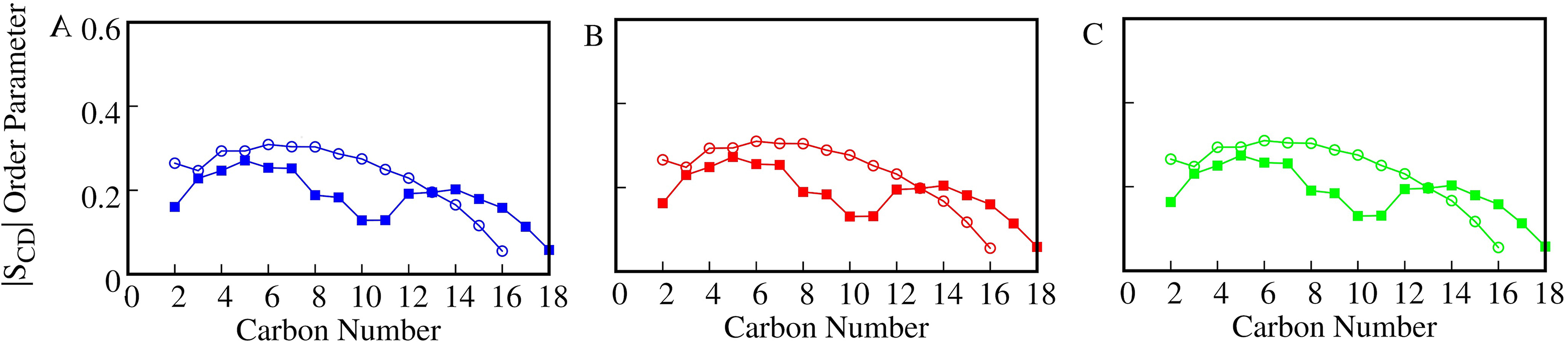
Order Parameter (S_cd_) as function of the position of the carbon atoms. The panels A, B and C correspond to the Scd of control, Withaferin-A and Withanone, in sn-1(-**∘**-) and sn-2 (-◼-) hydrocarbon chains.

The calculated order parameter shows the acceptable agreement between the experimental and calculated values. However, the small differences can be attributed to the inherent error of theoretical methods applied for the derivation of SCD values from NMR spectroscopy and simulations.

#### Potential of Mean Force (PMF)

In the solubility-diffusion model, the calculation of PMF is a critical component for the estimation of membrane permeability. It corresponds to the relative solubility of a permeant in solution *vs.* the membrane interior. The respective permeability of Wi-A and Wi-N from water phase to the interior of membrane is shown in Fig. 5.

**Fig. 5:**
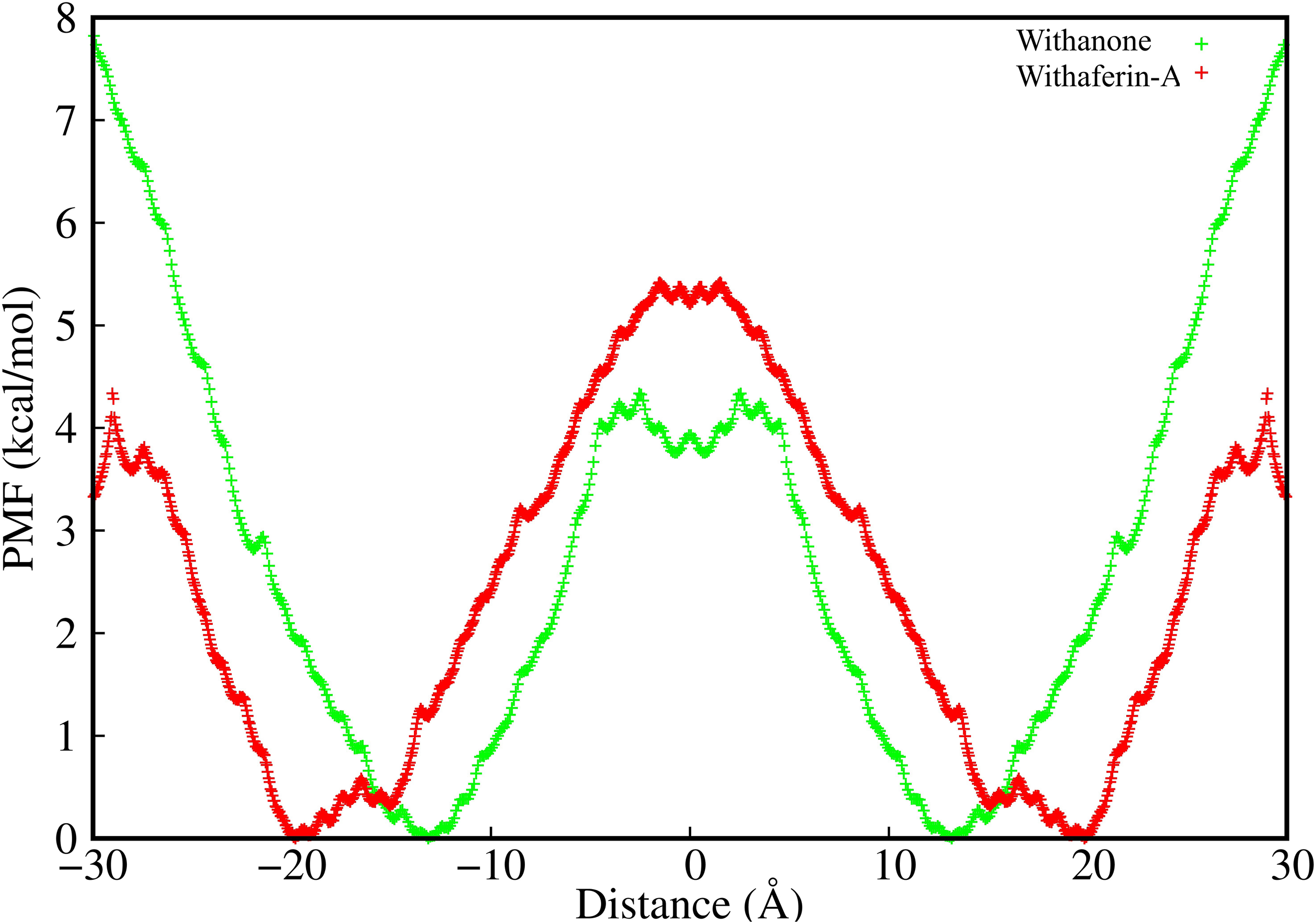
Change in the free energy (kcal/mol) of Withaferin-A (red) and Withanone (green) across the POPC bilayer.

Considering the first part of the curves, from bulk water region to the polar head group, one can identify a clear increase in the free energy for Wi-N as compared to Wi-A. This suggests a high permeability barrier for Wi-N in this region. Proceeding deeper in the bilayer that corresponds to the glycerol region, both the drug candidates display a free energy decrease, with a deep free energy minima at ∼20Å; therefore, this region can be considered as the preferably partitioned region. The rightmost part of the Fig. 5 shows the change in free energy characterizing the permeants in the hydrocarbon tail region (bilayer core). Here, both drugs showed an increase in free energy, highlighting the barrier effect inserted by the lipid hydrocarbon tail. However, the differences in free energy values are small for both drug candidates.

Further, the increase in free energy value for all solutes moving from bulk water phase into the membrane is generally attributed to the increase in density (Fig. 3). This lowers the free volume available to locate a permeant molecule and hence makes the solubilization of a solute more difficult. For the hydrophilic molecules, the lipid tail represents the main barrier to permeation. For hydrophobic molecules, partition is more favored in the middle of a bilayer than at lipid/water interface.

logP is the measure of lipophilicity of a molecule in un-ionized water, while logD is the measure of the same in ionized environment. As majority of known drugs are likely to be charged at physiological pH, using logP to describe their lipophilicity is misleading. logP only describes the partition coefficient of a neutral (uncharged) molecule hence logP offers an advantage in such (neutral stage) cases. The difference in the value of logP and logD can also be used as a measurement of understanding the effect of charge distribution on the surface of molecule. POPC membrane being highly chemically diverse, its hydrophilic layers provided charged environment to the incoming molecules. The logP value of Wi-A was slightly higher than that of Wi-N, but logD value of Wi-A was much higher than that of Wi-N. Besides, the interior of lipid membrane favors the transport of lipophilic molecules. This suggests that in charged environment, Wi-A is more lipophilic than Wi-N and hence this charge distribution over the surface of Wi-A and Wi-N could explain the selective permeability of the Wi-N. A video showing the permeation of Wi-A across the normal bilayer is included in the supplementary material (**Supplementary Video 1: Wi-A_premeability.mov**).

#### Resistivity

The second key component of the solubility-diffusion model is the diffusivity/Resistivity. The above results have indicated that transfer of Wi-A across the bilayer is less energy-driven and hence favorable as compared to Wi-N. Further, to check the resistivity exhibited by the bilayer on the drug molecules, the resistance exerted by the bilayer on both the natural drug molecules were calculated and shown in Fig. 6. The variation in resistivity showed a high resistance for Wi-N molecule in the head group region, while the resistance shown the bilayer is almost similar for both Wi-A and Wi-N in the tail region.

**Fig. 6:**
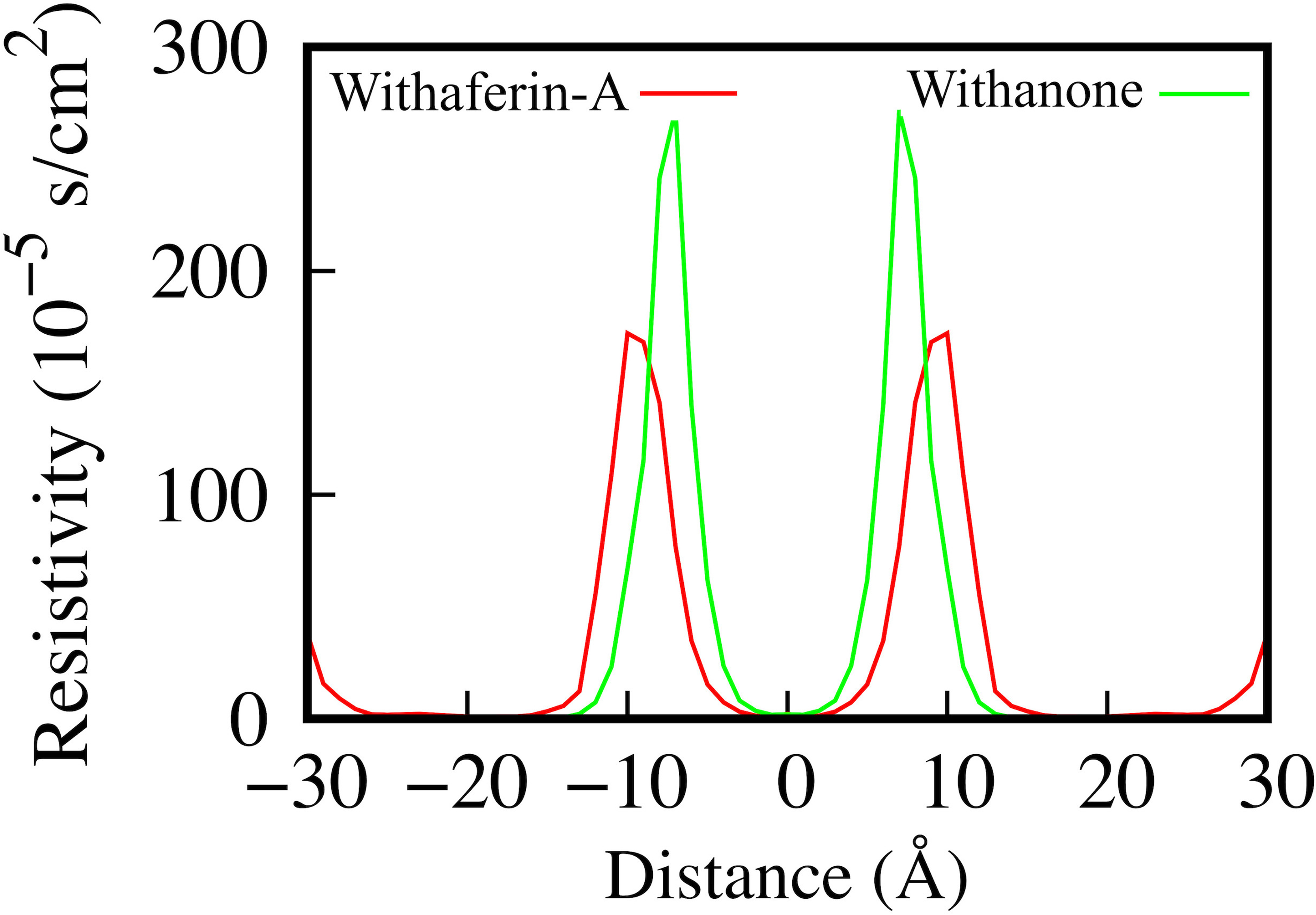
Resistivity profiles of Withaferin-A (red) and Withanone (green) molecules across the bilayer.

From the PMF (Fig. 5) and resistivity variation (Fig. 6), it is evident that the polar part of the membrane was primarily resisting the passage of natural drug molecule across the bilayer, while the hydrophobic tail region was offering almost similar barrier/resistance for both the withanolides.

#### Solvation and Hydrogen Bond

Though both the withanolides possess similar structure, our results showed a favorable passage for Wi-A across the bilayer. To understand the differences associated with the phenomenon, the average numbers of water molecules in the first hydration shell (number of water molecules in the 3.5Å radius of oxygen atom) were calculated for the drug candidates. The boxplot representation showed a higher solvation of Wi-A as compared to Wi-N, where the average solvation of epoxy oxygen is similar for both the permeants (Fig. 7). Though subtle differences could be observed for other oxygen atoms. In Wi-N, the average water solvation was ∼6 water molecules for rest of five oxygen atoms. However, for Wi-A, all the oxygen atoms were more solvated with ∼7 water molecules.

**Fig. 7:**
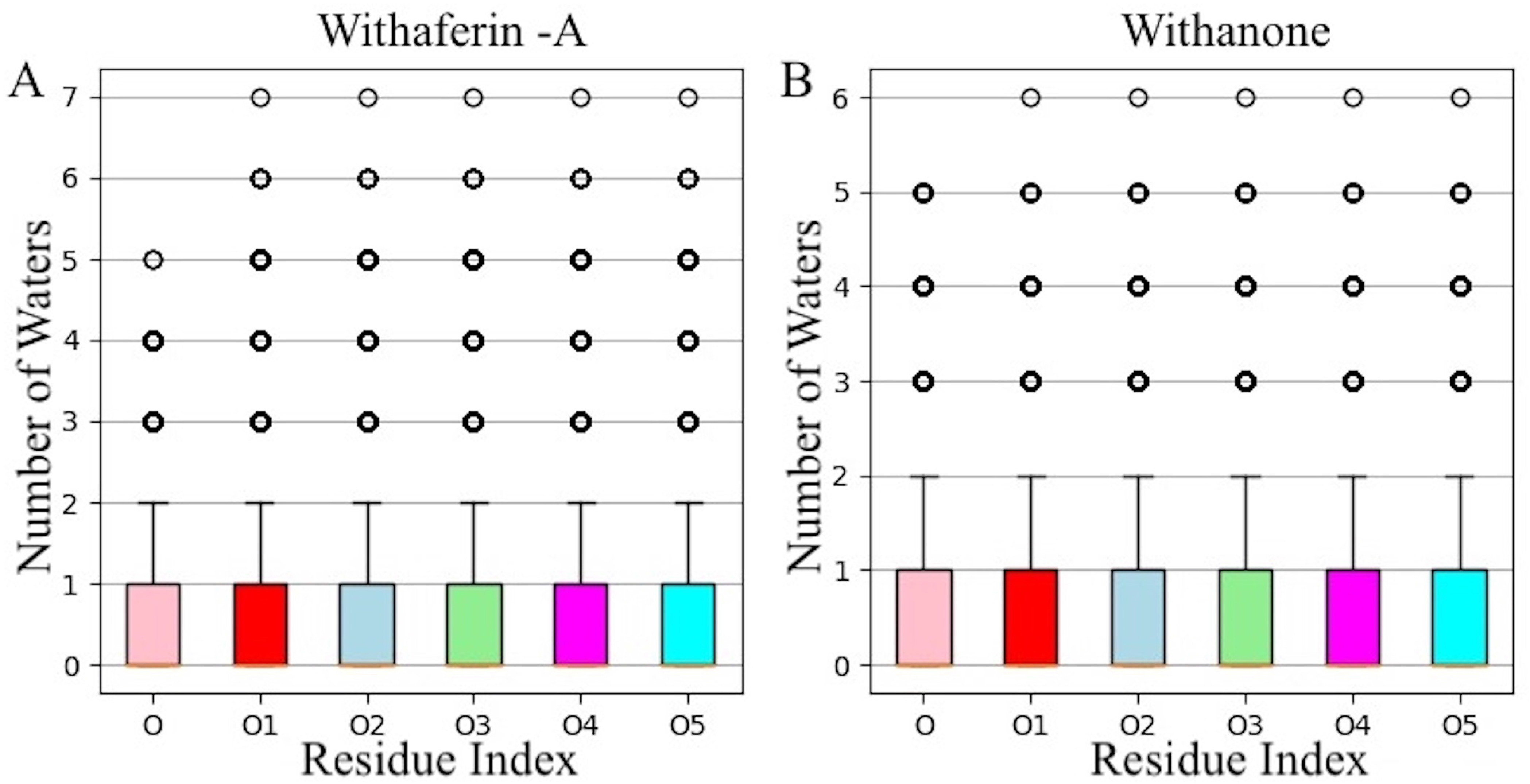
The average number of water molecules in the first hydration shell of oxygen atoms of Withaferin-A (A) and Withanone (B) molecules. The indexes of oxygen atoms are as mentioned in Fig.1.

#### Hydrogen Bond Lifetime

To get a clear picture of water dynamics around the drug molecules, the hydrogen bond lifetime between water molecule and drug oxygen atoms were investigated. Oxygen of water has a natural tendency to form hydrogen bond with other electronegative atom and hydrogen atom of water has the similar tendency to form hydrogen bond. Both the drug molecules have several –OH group that can form hydrogen bonds with water molecules; both water and drug molecules could act as hydrogen bond donor and acceptor in the formation of hydrogen bond. Depending upon the accessibility as well as the ambient environment, the interaction between the water and drug molecules may differ. Therefore, the calculation of hydrogen bond lifetime was expected to provide a measure for the elucidation of water dynamics around drug molecules leading to the favorable passage of Wi-A when compared to Wi-N. The maximum lifetime of hydrogen bond formed between water the terminal O4 and O5 oxygen atom of the Wi-A as compared to other oxygen atoms is shown in Fig. 8. The average maximum lifetime (in pico seconds) was found to be ∼5.10 ps and ∼4.98 ps respectively for O5 and O4 atoms of Wi-A. However, in case of Wi-N the average maximum lifetime was found to be ∼3.69 ps and ∼3.27 ps respectively for O5 and O4 atoms.

**Fig. 8:**
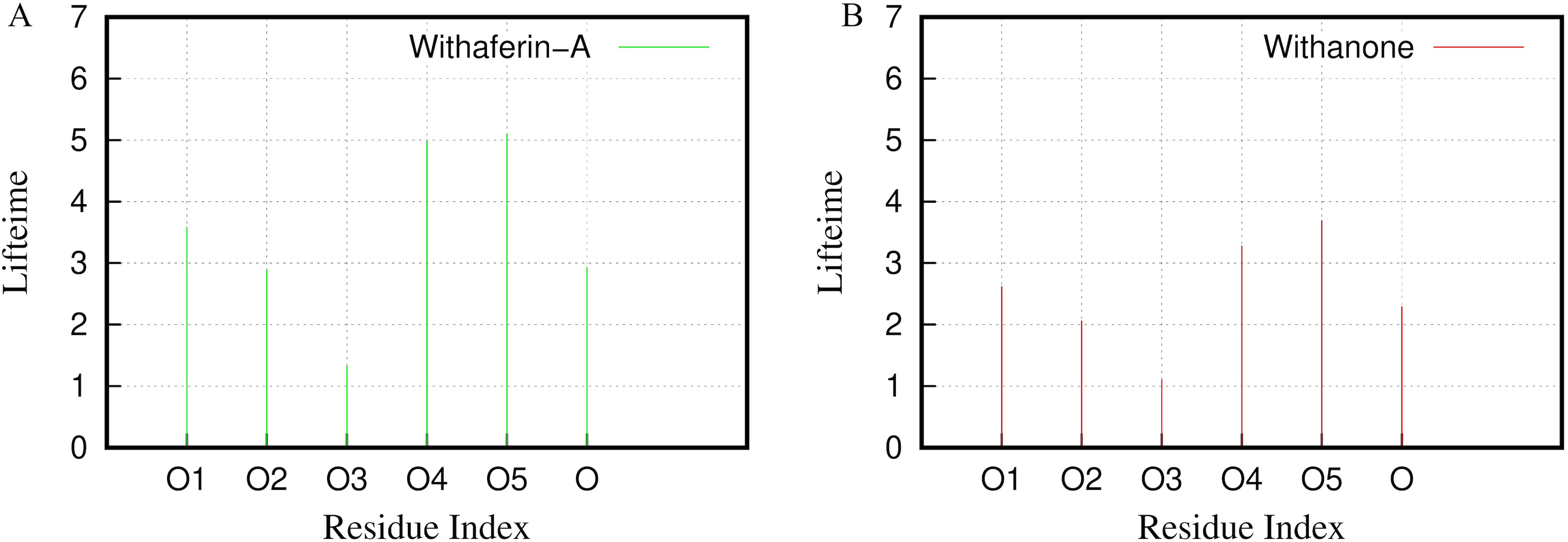
Maximum Hydrogen Bond Lifetime (in picosecond) between the oxygen atoms of Withaferin-A (A) and Withanone (B) drug molecules. Residue indexes for the oxygen atoms are as mentioned in Fig. 1.

#### Radial Distribution Function (RDF)

This analysis was carried out to track the passage of Wi-A molecule, which forms hydrogen bond with water, phosphate and oxygen atoms of the POPC atoms. The RDF between the oxygen atoms of Wi-A and lipid phosphate (Fig. 9A) and lipid nitrogen (Fig. 9B) was plotted. The nearest neighbor peak was clearly visible at ∼3.5Å and 4Å for the phosphorus and nitrogen groups respectively. In case of phosphorus, the second nearest-neighbor peak was also pronounced at ∼5Å. Among all the oxygen atoms of Wi-A, a strong interaction with lipid phosphate was observed with O5 followed by O4 and O1 atoms. Similarly, the O5 and O4 atoms again showed a high interaction with the lipid nitrogen atom. In case of Wi-N, the epoxy oxygen seemed to have a tight interaction with the lipid phosphorus and nitrogen atoms; however, the strength of this interaction was small as compared to Wi-A. Though both the molecules possess similar structure, still Wi-A cross the bilayer proficiently but not Wi-N. The difference between two molecules actually lies in the distribution of single hydroxyl group. In Wi-A, the hydroxyl group is attached to the terminal lactone ring, whereas in Wi-N, hydroxyl group is attached to the middle of steroidal ring. Though the number of total polar atoms in Wi-A and Wi-N remains the same, the position of polar atoms makes a big difference in the charge distribution over their surface (Fig. 1). The terminal arrangement of polar atoms in Wi-A provided it an advantage over the distributed polar atoms in Wi-N. Grouping of the polar atoms at just the terminals made the middle surface of Wi-A continuously non-polar favoring interactions with both hydrophilic and hydrophobic environment. Grouped polar atoms are more capable to face the hydrophobic environment as their own interactions can reduce the total exposed surface area.

**Fig. 9:**
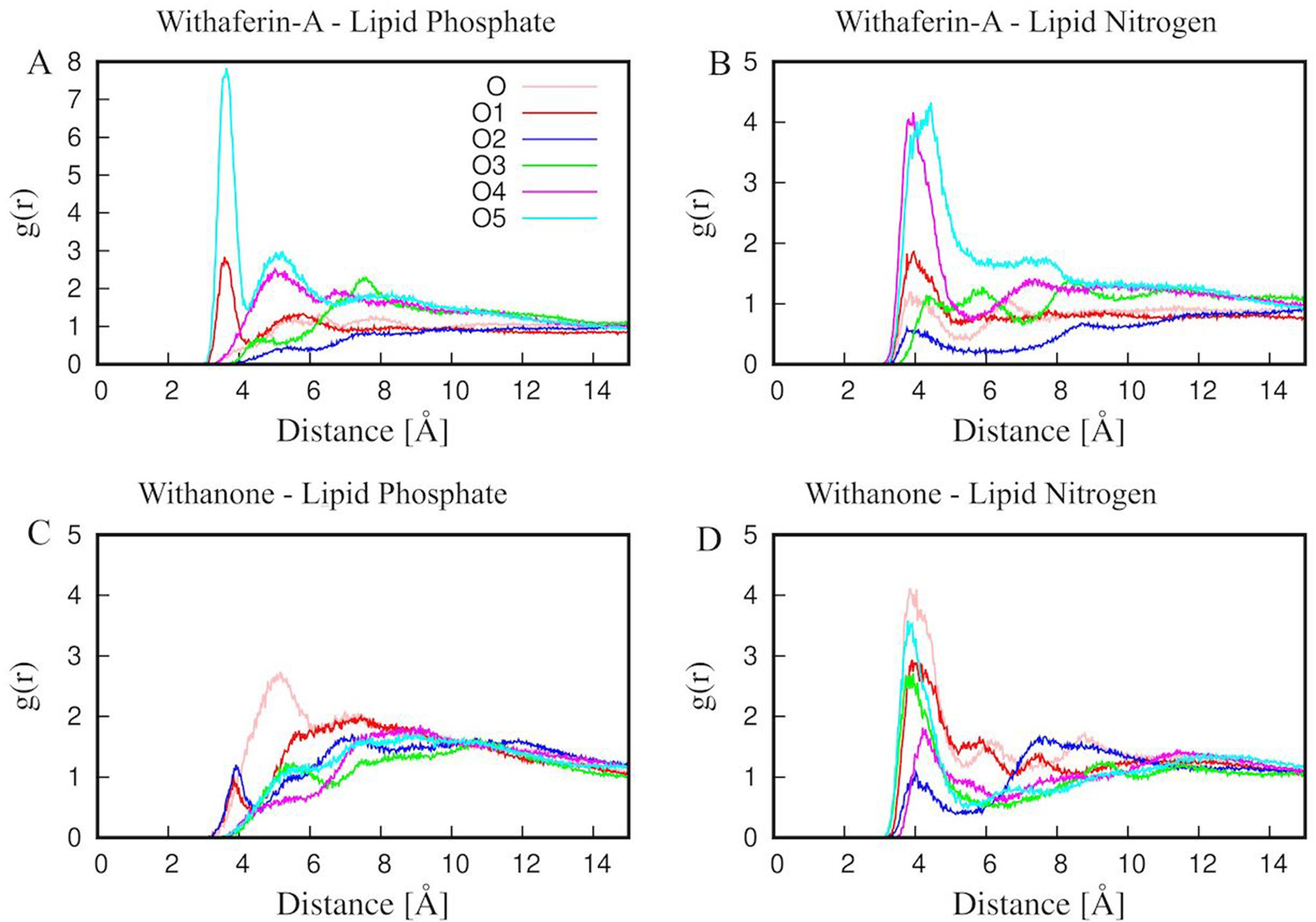
Radial Distribution Function (RDF) between oxygen atoms of Withaferin-A and Withanone with the lipid phosphate (Panels A and C) and nitrogen (Panels B and D) atoms. The index and color code for each oxygen is as mentioned in the top right of Panel A.

From the high solvation (Fig. 7) of terminal hydroxyl groups (O4 and O5) in Wi-A as well as the strong hydrogen bond lifetime (Fig. 8) of same atoms along with the RDF analysis (Fig. 9), it can be interpreted that the high interaction of terminal O5 and O4 atoms of Wi-A with polar atoms (water, lipid nitrogen, and lipid phosphate) of system enables the smooth transverse of Withaferin-A. While for Withanone, no such favorable interactions are found; hence the passage of Wi-N across the normal membrane is associated with high-energy cost.

### Experimental evidence to the differential absorption of Wi-A and Wi-N by normal cells

In order to provide the experimental evidence to the above membrane permeability analysis, we generated, for the first time, antibodies to Wi-A and Wi-N. As shown in **Supplementary Fig. 1**, 3/96 clones that initially showed reactivity to Wi-A and Wi-N were developed into hybridoma and were examined for their reactivity to Wi-A and/or Wi-N. We found that the one clone (L7C3-6) was able to detect Wi-A and Wi-N in cells treated with either purified compounds or i-Extract containing Wi-A and Wi-N as major components. The clone was designated as WiNA antibody. We next used anti-WiNA antibody to detect Wi-A/Wi-N in human cancer cells treated with purified Wi-A or Wi-N; isotype matched secondary antibodies were used as negative control. As shown in Fig. 10A, higher level of both Wi-A and Wi-N were detected in cells treated for 48 h as compared to 24 h. We extended the analysis to human normal cells. Wi-A, but not Wi-N was detected in the normal cells incubated with the purified compounds for 48 h (Fig. 10B). Furthermore, we found that Wi-A and Wi-N staining on the nucleus, which was confirmed by confocal microscopy (Fig. 10C). Wi-A was detected clearly in the nucleus of cancer as well as normal cells. In contrast, Wi-N was detected in the nucleus of cancer cells only (Fig. 10C). In order to further support these findings, we examined the expression of several proteins that have been shown to be involved in anticancer activity of Wi-A and Wi-N. As expected, Wi-A caused stronger up-regulation of γH2AX and p53, markers of DNA damage response in cancer cells (shown in Fig. 10D and E). Furthermore, only the cells treated with Wi-A showed decrease in CARF protein signifying apoptosis (Fig. 10E). The same was not observed in Wi-N treated cells. In light of these findings that endorsed reliable detection of Wi-A/Wi-N in cells with our antibodies, we expanded the analysis in more human normal cells. We found that Wi-A entered the cells and triggered DNA damage response in normal cells as determined by increase in H2AX, pATR and pCHK1 (Fig. 10F and G). In contrast, Wi-N induced increase in these marker proteins was not detected in normal cells. These data confirmed the MD predictions for permeability of Wi-A and Wi-N through normal cell membrane, and for the first time demonstrate that the selective toxicity of Wi-N to cancer cells is mediated by its poor capability to enter normal cells.

**Fig. 10:**
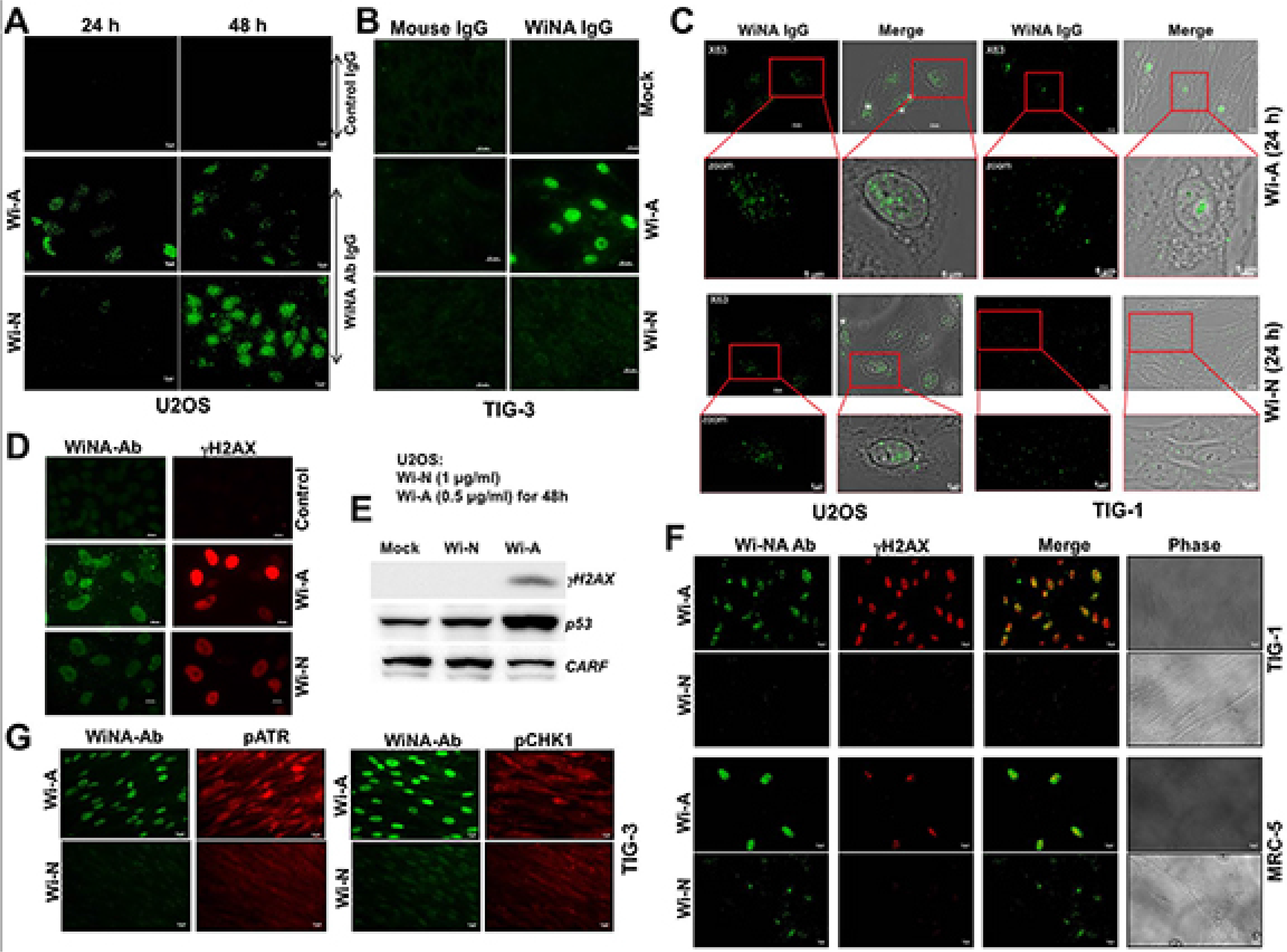
Experimental validation of differential permeation of Withaferin-A (Wi-A) and Withanone (Wi-N) through cell membrane. **(A)** Human cancer cells (U2OS) treated with Wi-A/Wi-N that were immunostained with anti-WiNA Ab showed their presence in cells. (**B**) Normal cells treated with Wi-A/Wi-N for 48h showed presence of Wi-A, but not Wi-N. (**C**) high resolution images showing the presence of Wi-A in U2OS as well as TIG-3 cells. Wi-N was detected only on U2OS cells. (**D**) Wi-A/Wi-N treated cells were examined for DNA damage marker protein γH2AX showed its higher upregulation and activation in Wi-A, as compared to Wi-N treated U2OS cells. (**E**) Upregulation of γH2AX, p53 and down regulation of CARF in Wi-A, but not Wi-N, treated cells. (**F and G**) Wi-A, but not Wi-N, treated normal (TIG-3 and MRC5) cells showed increase in γH2AX, pATR and pCHK1.

## Conclusion

To the best of our knowledge, this is the first detailed computational study characterizing the permeation of two anti-cancer natural drugs (withanolides) across the lipid bilayer. We have used computational as well as experimental methods to predict the permeability of drug compounds. Taking together all the data, we observed that the permeation of Wi-A and Wi-N doesn’t affect the membrane integrity. The values obtained by area per lipid, electron density and order parameters were almost similar for bilayer with and without these drug molecules. The PMF calculation showed that the permeation of Wi-A across the bilayer was associated with low energy barrier and therefore, could be considered as a favorable transverse. The free energy barrier was large for Wi-N and hence had an unfavorable passage across the bilayer. Further, these findings were confirmed by the resistivity profiles of both the natural drug molecules. Wi-N experienced a high resistance when compared to Wi-A. To elucidate the mechanics for this behavior, the solvation and hydrogen bond analysis was carried out, which showed clearly that the high solvation of terminal hydroxyl group (O5) in the Wi-A provides a favorable interaction of Wi-A with the polar head group of the membrane, thus facilitating a smooth passage for Wi-A. These results were further confirmed by the radial distribution function. It was observed that the strong interaction between Wi-A-O5 with polar phosphate/nitrogen atoms of lipid bilayer was permitting the smooth passage for this drug. Nonetheless, the logP and logD values associated with the Wi-A and Wi-N supported our results. In spite of the fact that Wi-N also has similar structure like Wi-A, the lipophilicity of the molecule was less as compared to Wi-A. Thus, the passage of Wi-N exhibits a high resistance. The difference in the permeation between Wi-A and Wi-N could therefore be attributed to the difference in the position of hydroxyl group on steroidal lactone ring. This study has revealed the fact that the subtle difference associated with the positioning of functional groups can play a major role in transportation of drug molecules across the bilayer. Our experimental data also corroborates the computational findings. The experimental data demonstrated that Wi-A, but not Wi-N, proficiently entered the normal cells that were tracked with unique withanolide-recognizing antibodies. Since cancer cells are actively dividing and possess high energy than normal cells, Wi-N could permeate cancer, but not normal cells. On the other hand, Wi-A, due to low energy barrier, traverses through both cancer and normal cell membranes. This may explain, selective toxicity of Wi-N to cancer cells, at least in part. Although the calculated free energy profile for the permeation of Wi-A and Wi-N were qualitatively matching with the experimental observation, explanation for adequate effect of drug passage on the membrane permeability requires additional simulation with varying drug concentration. This would shed further light on the proposed mechanism. Our work is an important step towards understanding the molecular basis of permeability of natural drug molecules. More membrane models can be generated with varying composition and density, representing different normal and cancer cell types, and can be used for similar permeability studies for small molecular compounds.

## Acknowledgements

We acknowledge the High-Performance Computing (HPC) Facility of IIT Delhi and the Bioinformatics Centre supported by the Department of Biotechnology (Govt. of India) at IIT Delhi, for computational resources. This study was made possible in part through the support of Department of Biotechnology (Government of India) for DAILAB at IIT Delhi (India) and AIST (Japan).

## Author contributions

NSY and SPK designed the model and the computational framework for this study. TY, CL and HA did the experiments. RW, SCK, CY and DS conceived the study and wrote the paper with inputs from all the authors.

